# Epigenetic influences on aging: a longitudinal genome-wide methylation study in old Swedish twins

**DOI:** 10.1101/226266

**Authors:** Yunzhang Wang, Robert Karlsson, Erik Lampa, Qian Zhang, Åsa K. Hedman, Malin Almgren, Catarina Almqvist, Allan F. McRae, Riccardo Marioni, Erik Ingelsson, Peter M. Visscher, Ian J. Deary, Lars Lind, Tiffany Morris, Stephan Beck, Nancy L. Pedersen, Sara Hägg

## Abstract

Age-related changes in DNA methylation have been observed in many cross-sectional studies, but longitudinal evidence is still very limited. Here, we aimed to characterize longitudinal age-related methylation patterns (Illumina HumanMethylation450 array) using 1011 blood samples collected from 385 old Swedish twins (mean age of 69 at baseline) up to five times over 20 years. We identified 1316 age-associated methylation sites (p<1.3×10^−7^) using a longitudinal epigenome-wide association study design. We measured how estimated cellular compositions changed with age and how much they confounded the age effect. We validated the results in two independent longitudinal cohorts, where 118 CpGs were replicated in PIVUS (p<3.9×10^−5^) and 594 were replicated in LBC (p<5.1×10^−5^). Functional annotation of age-associated CpGs showed enrichment in CCCTC-binding factor (CTCF) and other unannotated transcription factor binding sites. We further investigated genetic influences on methylation (methylation quantitative trait loci) and found no interaction between age and genetic effects in the 1316 age-associated CpGs. Moreover, in the same CpGs, methylation differences within twin pairs increased over time, where monozygotic twins had smaller intra-pair differences than dizygotic twins. We show that age-related methylation changes persist in a longitudinal perspective, and are fairly stable across cohorts. Moreover, the changes are under genetic influence, although this effect is independent of age. In addition, inter-individual methylation variations increase over time, especially in age-associated CpGs, indicating the increase of environmental contributions on DNA methylation with age.

## Introduction

DNA methylation is known as a key factor in human aging (1). In the human aging process, alterations of DNA methylation indicate a loss of epigenetic control and relate to pathological phenotypes (2, 3). Today, epigenome-wide association studies (EWAS) have established general knowledge on age-related methylation patterns in humans (4–8). Overall, approximately 30% of cytosine-phosphate-guanine (CpG) sites measured by Illumina 450k array are associated with age (7, 9). and they can be divided into hypo- or hypermethylations (10). Moreover, individual variation of DNA methylation partly depends on genetic variants and methylation is believed to be a mediator of genetic effects (11). Several studies on methylation quantitative trait loci (meQTL) have identified genetic associations with methylation (2, 12). In addition, it has been suggested that age-related methylation alterations depend on genetic effects (9), and another recent study on mother-children pairs reported that meQTL associations were stable over time (12). Twin studies of the epigenome have been used to estimate methylation heritability (9, 13), and higher methylation correlation has been observed in monozygotic (MZ) than dizygotic (DZ) twin pairs (5, 14). However, the change of intra-twin-pair methylation difference over time has not been studied.

To date, most EWAS publications on age used cross-sectional data, while longitudinal studies on intra-individual change in methylation over time are still sparse (15). Hence, the aim of this study was to investigate longitudinal age-related alterations in DNA methylation, using whole blood samples from old Swedish twins (mean age of 69 at baseline) collected up to five times across 20 years. Age-associated CpGs were then validated in two independent longitudinal cohorts. Age-related changes of cellular compositions were estimated and their effects on age-related change of methylation were measured. In addition, we analyzed meQTL associations and studied genetic effects on age-related methylation patterns over time, with specific emphasis on twin-pair differences.

## Results

### Longitudinal EWAS on age

Using the Illumina HumanMethylation450k array, DNA methylation data of 390,894 autosomal CpGs were obtained from 385 twins (73 MZ, 96 DZ complete twin pairs) enrolled in the Swedish Adoption/Twin Study of Aging (SATSA) (16) (Table 1), which is part of the Swedish Twin Registry (17). The longitudinal EWAS analysis revealed systematic changes of methylation with age. In total 1316 CpGs were identified as significantly associated with age using a Bonferroni-corrected threshold (p<1.3×10^−7^) (Figure S1, File S1), where 1026 CpGs were hypomethylated. The top CpG was found within the gene *ELOVL2* (Figure S2). Sex was adjusted for as a covariate in the model, and 6509 CpGs were significantly associated with sex (p<1.3×10^−7^), of which 16 were also found among the 1316 age-related CpGs. Overall, sex-associated CpGs were not enriched in the age-associated CpGs. A sensitivity analysis showed that twin zygosity had little impact on results.

**Table 1.** Characteristics of the longitudinal DNA methylation samples collection in SATSA.

| Longitudinal wave | Year of sample collection | Number of Participants (new recruits)* | Female Proportion | Age mean (SD) |
| --- | --- | --- | --- | --- |
| 1 | 1992-1994 | 239 | 59% | 68.6 (9.1) |
| 2 | 1999-2001 | 242 (102) | 63% | 71.2 (10.1) |
| 3 | 2002-2004 | 188 (26) | 54% | 72.1 (9.1) |
| 4 | 2008-2010 | 186 (15) | 61% | 76.2 (8.5) |
| 5 | 2010-2012 | 156 (3) | 66% | 77.9 (8.4) |

\* The numbers of newly recruited participants in each wave in parenthesis. 200 participants were available in at least three waves. SATSA, the Swedish Adoption/Twin Study of Aging; SD, standard deviation.

Validation of age-associated CpGs was done in two independent longitudinal cohorts; the Prospective Investigation of the Vasculature in Uppsala Seniors (PIVUS) (18) and the Lothian Birth Cohort (LBC) (19). The PIVUS cohort measured methylation data from 196 individuals at age 70 and 80, and the LBC cohort, including two sub-cohorts of 906 and 436 individuals at baseline, measured blood samples at three time points, with mean ages of 70 and 79 at baseline (Table S1). Among the 1316 age-associated CpGs identified in SATSA, 1271 and 973 CpGs were available in PIVUS and LBC respectively. In PIVUS, 118 of the 1271 CpGs were consistent in effect directions and significant at a Bonferroni-correction threshold for validation p<3.9×10^−5^. In LBC, 594 out of 973 CpGs were consistent in effect directions and significant with p<5.1×10^−5^. The correlation of effect sizes between PIVUS and SATSA was 0.57, and 0.87 between LBC and SATSA (Figure 1).

**Figure 1.**
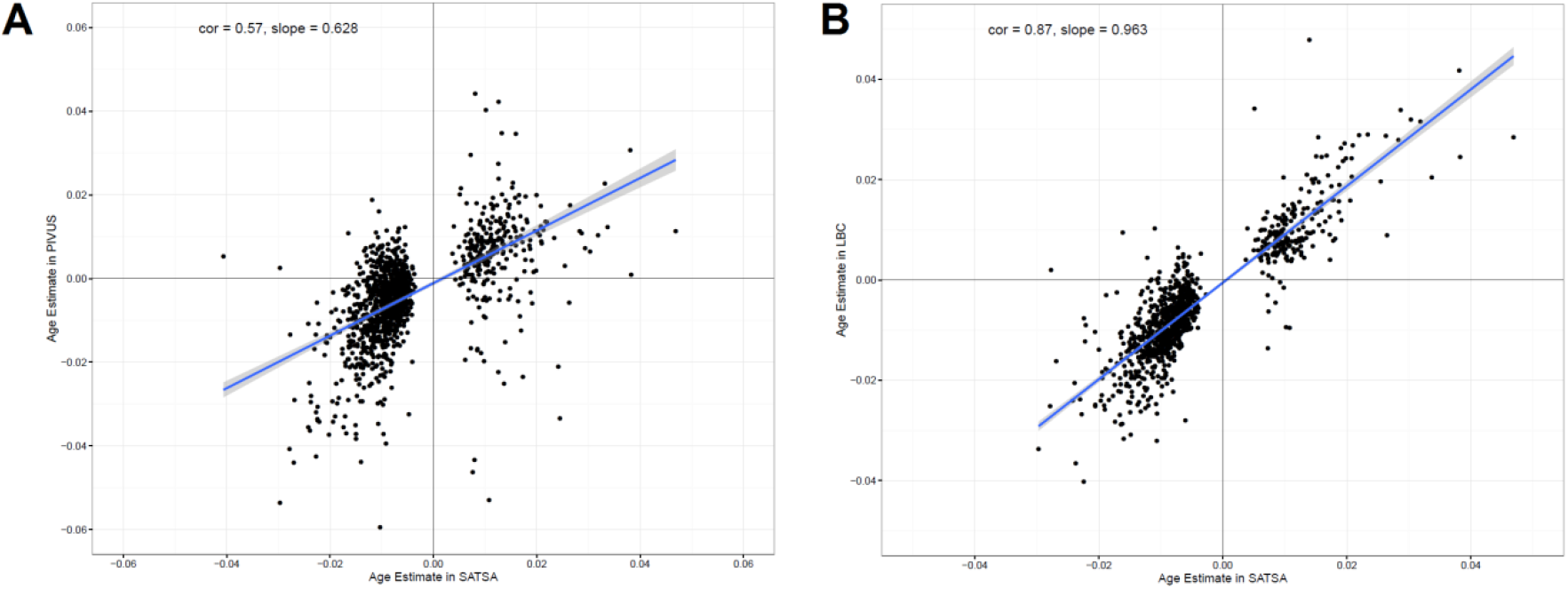
The effect sizes of age on age-associated CpGs in SATSA and two independent longitudinal cohorts. Effect sizes of age were estimated from a longitudinal epigenome-wide associated study of age in SATSA, using a mixed effect model. The 1316 Bonferroni significant CpGs (p<1.3×10^−7^) were tested for age associations in PIVUS and LBC. A) The Pearson correlation of the effect sizes is 0.57 (p<10^−16^) between PIVUS and SATSA. The slope of the linear regression line is 0.63. B) The Pearson correlation is 0.87 (p<10^−16^) between SATSA and LBC. The slope of linear regression is 0.96.

### Cellular compositions change with age

Cellular compositions were estimated from methylation data using the method by Houseman (20) and the longitudinal change of cellular compositions with age was measured using a mixed effect model. The total peripheral blood mononuclear cell (PBMC) proportions increased with age (p=4.6×10^−8^) while granulocytes proportions decreased with age (p=1.1×10^−4^). Within PBMCs, CD14+ monocytes (p=9.4×10^−16^) and natural killer cells (p=6.0×10^−4^) significantly increased with age, while CD19+ B cells decreased with age (p=9.2×10^−4^). Within granulocytes, eosinophils increased with age (p=2.6×10^−4^) while neutrophils did not change with age (Figure 2). However, in general, age only explains a small proportion of the variance of cellular compositions in our cohort (Figure 2, File S1).

**Figure 2.**
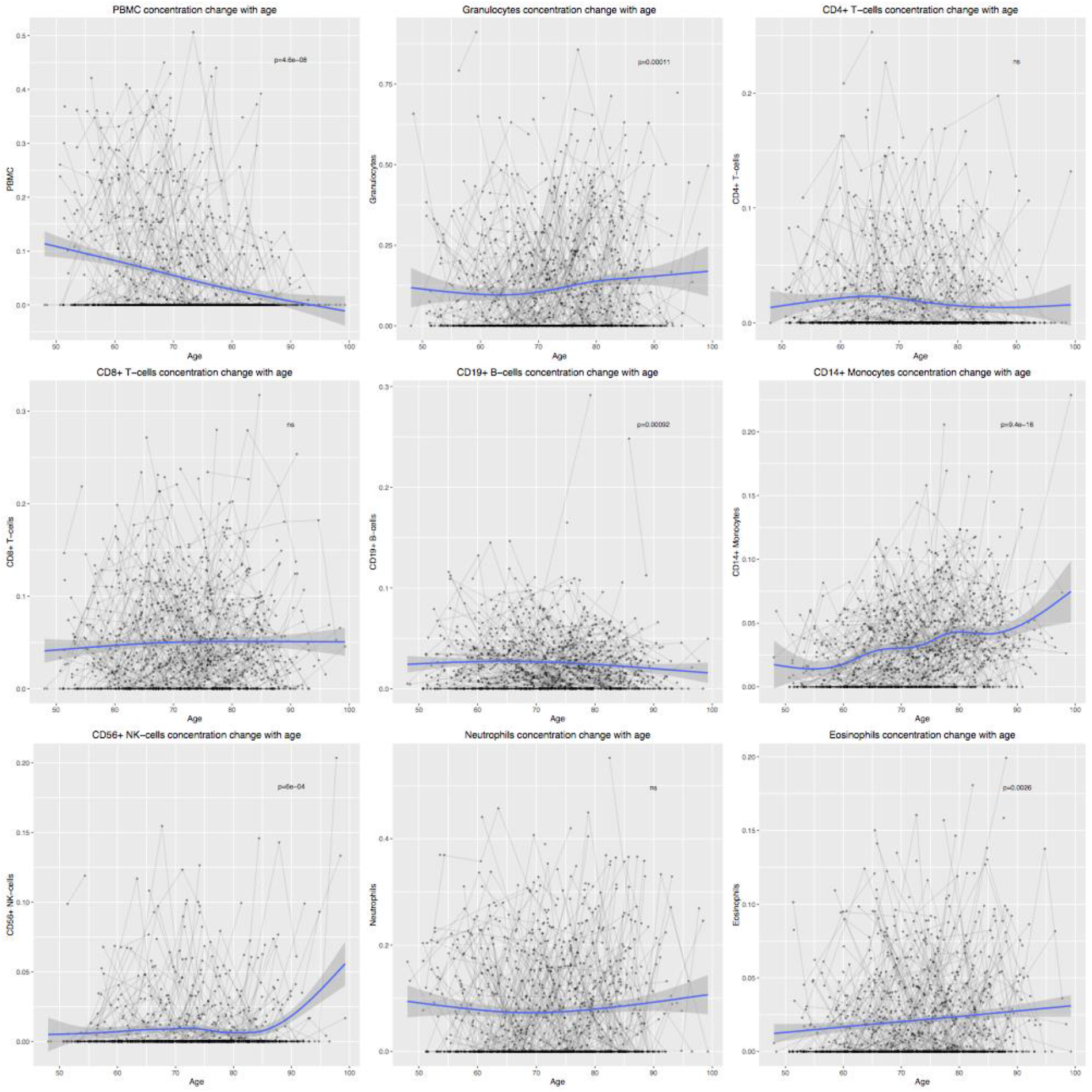
Longitudinal changes of cellular compositions with age. Estimated cellular compositions were plotted against age for each cell types. Grey lines indicate multiple observations of individuals. P-values were calculated from a mixed effect model measuring the longitudinal change of cellular proportions. PBMC, peripheral blood mononuclear cell; NK cell, natrual killer cell

Furthermore, the adjustment of cellular compositions in the 1316 age-associated CpGs only slightly increases the effect sizes of age, especially for hypermethylated CpGs. In total, 246 CpGs were significantly (p<3.8×10^−5^) associated with at least one cell type and 40 of them were associated with all estimated cell types. However, effect sizes of most CpGs did not change much (Figure S3, File S1).

### Regulatory and functional annotations of age-associated CpGs

The CpG island locations of the identified age-associated CpGs were obtained from the Illumina manifest file. Age-associated CpGs were less frequently found in CpG islands and open sea regions, and more frequently in CpG shores among the probes designed in the 450k chip. The majority (1026 of 1316) of the CpGs showed decreased methylation with age, and among the hypermethylated CpGs, the majority (85.2%) were located in CpG islands (Error! Reference source not found.A).

To explore biological functions, we annotated the age-associated CpGs using regulatory features from the Ensembl database, showing CpGs in relation with regulatory protein binding sites (21). Compared to regulatory features of the 450k probe background, age-associated CpGs were found enriched in CTCF binding sites, promoter flanking regions and other transcription factor binding sites (Figure 3B). In particular, a much higher proportion of hypermethylated CpGs were found in transcription factor binding sites than other regulatory features.

**Figure 1.**
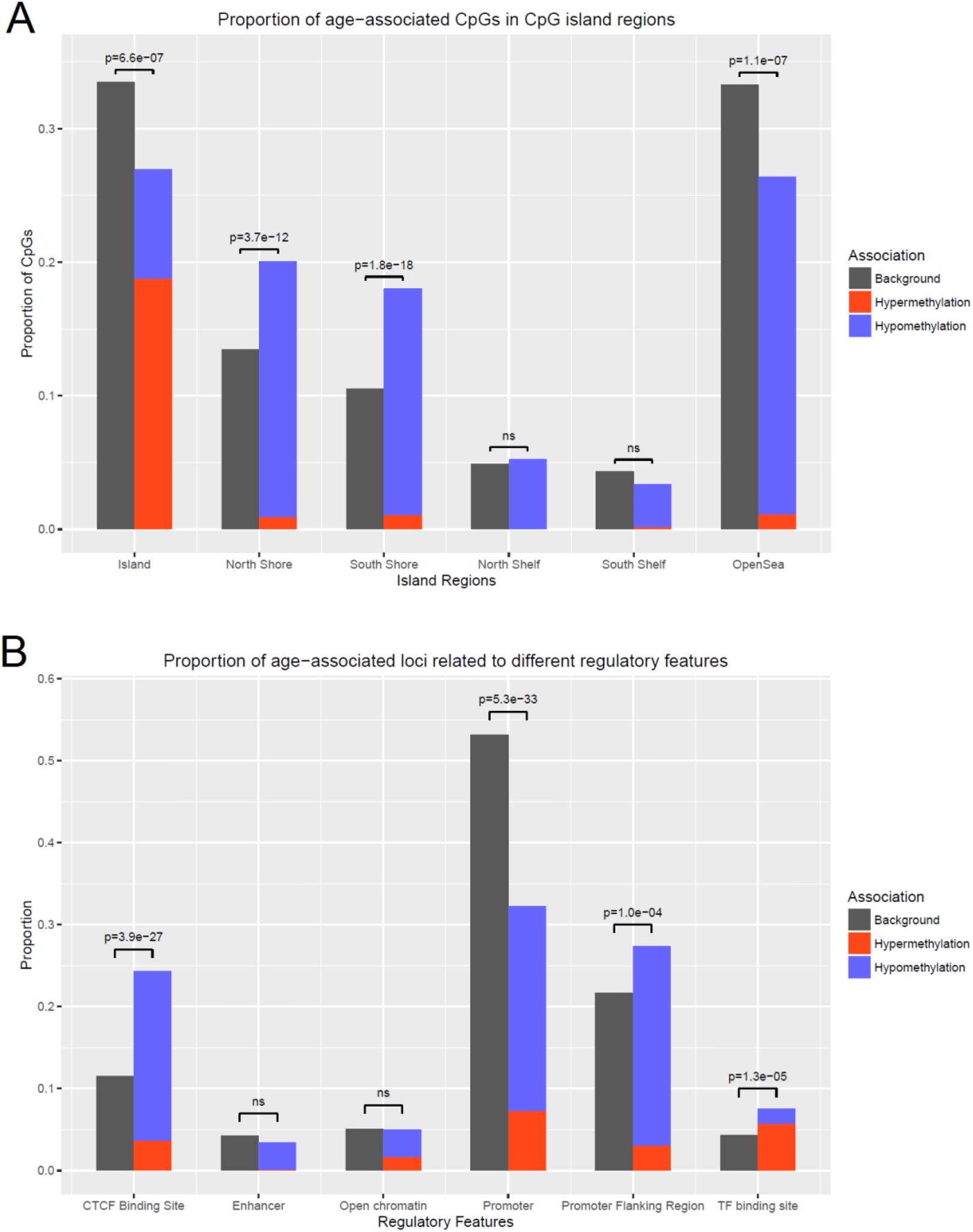
The distribution of age-associated CpGs in relation to CpG islands and regulatory features. A) Proportions of age-related hyper- and hypomethylated CpGs in different CpG island regions compared to proportions on the 450k array. Age-associated CpGs are enriched in CpG shores (North Shore p=3.7×10^−12^ and South Shore p=1.8×10^−18^), and depleted in CpG islands (p=6.6×10^−7^) and open sea regions (p=1.1×10^−17^). Outside of CpG islands, 918 out of 961 CpGs are hypomethylated with age, and in CpG islands, 247 out of 355 CpGs are hypermethylated with age. B) The proportions of age-related hyper- and hypomethylation, as well as background CpGs, in different regulatory regions. Age-associated CpGs are highly enriched in CTCF binding cites (p=3.9×10^−27^). Only in TF binding sites, the proportion of age-related hypermethylated CpGs is higher than hypomethylated CpGs. The enrichment or depletion is shown by p-values calculated from two-sample proportion tests. CpG, cytosine-phosphatate-guanine; ns, non-significant; CTCF, CCCTC-binding factor; TF, transcription factor.

We further mapped the 1316 age-associated CpGs to 878 genes according to the Illumina manifest file. Consequently, genes were annotated using the Database for Annotation, Visualization and Integrated Discovery (DAVID) online tool (22, 23), and 85.3% of the genes were categorized according to the biological process term of gene ontology (GO). Seven GO terms were found significantly enriched (false discovery rate [FDR] < 0.05), and the top function was homophilic cell adhesion via plasma membrane adhesion molecules (Table 2). Moreover, many of the genes were found to be enriched in functions related to nervous system development and neurogenesis.

**Table 2.** Enriched Gene Ontology terms for genes mapped to the age-associated CpGs.

| Gene Ontology term | Number of genes | p-value | FDR |
| --- | --- | --- | --- |
| Homophilic cell adhesion via plasma membrane adhesion molecules | 42 | $5.4 \times 10^{-22}$ | $1.0 \times 10^{-18}$ |
| Nervous system development | 147 | $1.3 \times 10^{-9}$ | $2.4 \times 10^{-6}$ |
| Neurogenesis | 100 | $6.0 \times 10^{-7}$ | $1.1 \times 10^{-3}$ |
| Organ morphogenesis | 73 | $1.2 \times 10^{-6}$ | $2.3 \times 10^{-3}$ |
| Cell development | 121 | $7.2 \times 10^{-6}$ | $1.3 \times 10^{-2}$ |
| Neuron differentiation | 83 | $1.9 \times 10^{-5}$ | $3.4 \times 10^{-2}$ |

CpG, cytosine-phosphatate-guanine; FDR, false discovery rate

### Identification of *cis*-meQTLs

To investigate genetic influences on methylation, *cis*-meQTLs (distance between markers <1 million base pairs) from 1.9 billion possible associations between 6.5 million single nucleotide polymorphisms (SNPs) and 390,894 CpGs across the genome were analyzed. Over 1.4 million associations were statistically significant using a Bonferroni-corrected threshold (p<2.5×10^−11^). As expected, we observed more associations and lower p-values when SNPs were closer to their associated CpGs (Figure S4). In total, 14,714 CpGs were significantly associated with at least one SNP.

Overall, our results were consistent with associations in the mQTL database (12). About 44% of SNP-CpG associations identified in SATSA were also significant (p<10^−14^) in the middle-age group in the mQTL database. Also, 8950 out of the 14,714 (61%) SNP-associated CpGs identified in SATSA were also significantly (p<10^−14^) associated with SNPs in the mQTL database.

### Genetic effects on age-associated CpGs

To investigate the relationship between age and genetic effects on methylation, we specifically studied CpGs that were significantly associated with both age and genetic variants. Among the 1316 age-associated CpGs discovered in the longitudinal analysis, 123 (9.3%) were also associated with at least one SNP. The proportion of genetic-associated CpGs among the age-associated CpGs (9.3%) was higher than the proportion in all CpGs (3.7%; p=6.7×10^−26^). For each of those CpGs, a longitudinal mixed effect model was performed including the associated SNP with the lowest p-value as a covariate. The age effects were still significant after adjusting for top associated SNPs (File S2). Moreover, interactions between age and SNPs were tested in the models as covariates, and no significant interaction was observed.

### Methylation differences within twin pairs

We further calculated standardized Euclidean distances from genome-wide methylation data to measure intra-pair differences between MZ and DZ twins over time (Table S3). Taking all CpGs into account, distances within twin pairs increased significantly with age (β=0.021, p=9.4×10^−4^) (Figure 4A), with steeper slopes when using the 1316 age-associated CpGs only (β=0.029, p=2.9×10^−5^; Figure 4B). The slope of the age effect on methylation differences in SNP-associated CpGs was smaller than that of all CpGs (β=0.015, p=3.32×10^−5^; Figure 4C).

**Figure 2.**
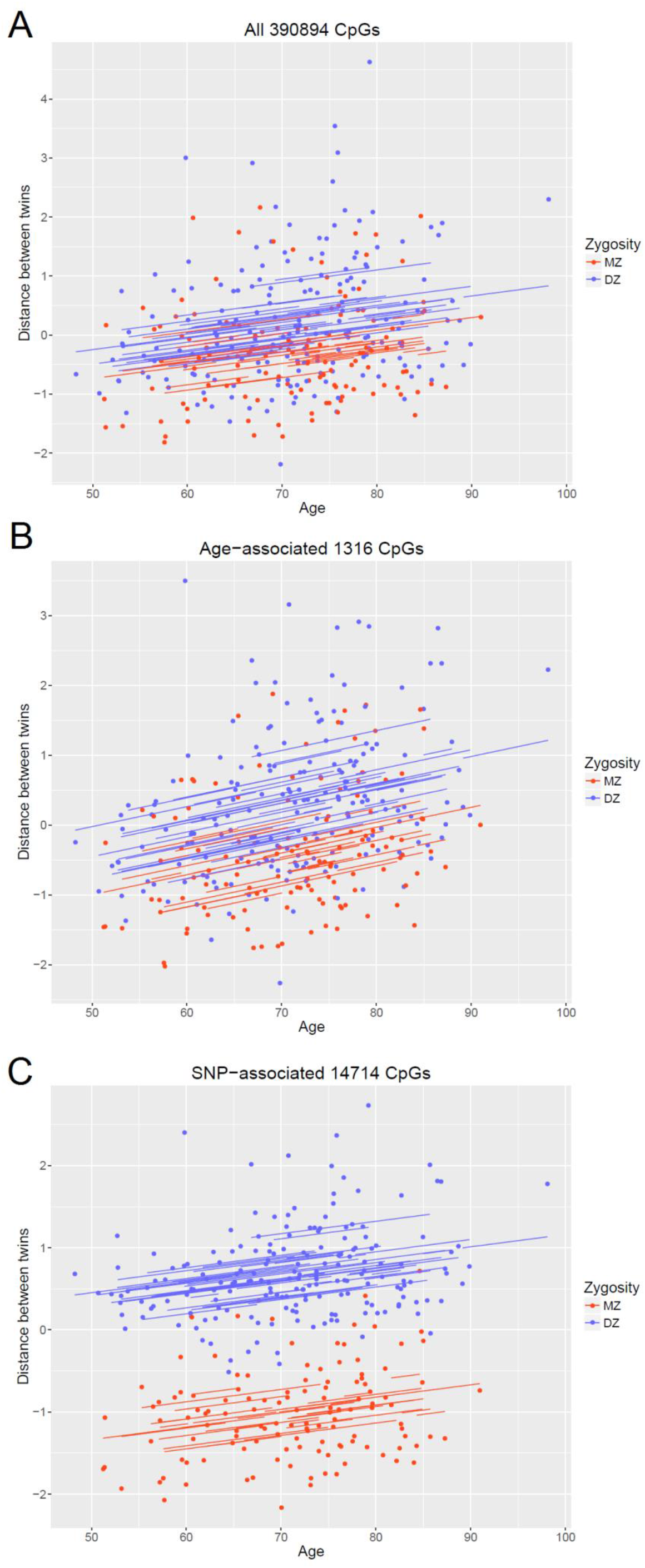
Regression plots of intra-twin-pair methylation differences over time in SATSA. Methylation differences within twin pairs at the same time point calculated by Euclidean distances of A) all CpGs, B) age-associated CpGs, and C) SNP-associated CpGs. Blue lines and points are MZ and red lines and points are DZ. Methylation differences within twin pairs increase with age, especially in age-associated CpGs. CpG, cytosine-phosphatate-guanine; MZ, monozygotic; DZ, dizygotic; SNP, single nucleotide polymorphism.

Furthermore, MZ and DZ twins showed significant intra-twin-pair Euclidean distances based on all CpGs, where MZ twins had significantly smaller distances compared to DZ twins (β=0.499, p=1.0×10^−4^; Figure 4A). In particular, the difference was much clearer for the 14714 SNP-associated CpGs (β=1.689, p=1.46×10^−65^; Figure 4C). The difference between zygosities for the 1316 age-associated CpGs was significant and the effect size is larger than that of all CpGs (β=0.697, p=6.75×10^−8^; Figure 4B).

Moreover, for each CpG, we performed an intra-twin-pair analysis to measure how much each one of them contributed to the total increasing methylation distances within twin pairs. Although no CpG passed the Bonferroni threshold (p<1e-7), almost all CpGs with p<0.05 have positive estimates of age, indicating that the differences in global methylation patterns between twins increased over time (Figure S5). On average, the age-associated CpGs have a higher effect size compared to all CpGs (15% higher, p=4.18×10^−05^ from a t-test), explaining the steeper slope of age-associated CpG in Figure 4. However, the CpGs that contributed to the growing intra-twin-pair methylation distances (p<0.05), have a much larger mean effect size than age-associated CpGs (Figure S5).

## Discussion

In this study, we characterized longitudinal age-related DNA methylation patterns in old twins, and identified 1316 CpGs associated with age. The strongest age-associated CpG was found in the promoter of the *ELOVL2* gene. Also, we described the longitudinal change of estimated cellular compositions with age and how cellular compositions affect age-associated methylation patterns. Moreover, genetic effects were observed for some age-associated CpGs, but they were independent of age effects. Furthermore, analyses of methylation differences within twin pairs revealed increasing differences over time, where MZ twins showed smaller dissimilarities than DZ twins in age-associated CpGs.

The age-related CpGs from SATSA were successfully validated in the longitudinal cohort LBC, but not so consistent in PIVUS. The LBC cohort validated 594 of the 1316 age-related CpGs at p<5.1×10^−5^, with a large sample size and three longitudinal waves. The high correlation of effect sizes of age (r=0.87) between SATSA and LBC indicated that LBC verified the effect sizes estimated by SATSA as well. The PIVUS cohort validated 118 of the 1316 CpGs at p<3.9×10^−5^ and had an intermediate correlation (r=0.57) of age effect sizes with SATSA. Specifically, PIVUS failed to validate the convincing top hit cg16867657 from SATSA. One explanation could be that the PIVUS cohort only collected data from participants at two ages, 70 and 80 years. Thus, the PIVUS cohort was quite different from SATSA in study design, where SATSA had up to five longitudinal waves and a much wider age range (49-99 year). Moreover, the age-related CpGs identified in this study were generally consistent with several published cross-sectional studies using 450k data (Table S). Johansson (7) and Dongen (9) reported around 30% of all CpGs to be age-related, which overlapped with a majority (91% and 66% respectively) of the 1316 age-associated CpGs we identified. Florath (8) only reported 162 significant CpGs and half were also found in SATSA. Our results did not overlap much with the longitudinal study by Tan (15), however, the set-up of the study with data from 43 twin pairs at two time points (10 years apart) was again different from ours. In general, when conducting a longitudinal EWAS study, many more covariates come into place and technical variation from different waves could bias the estimates. The SATSA methylation samples, however, were completely randomized on methylation arrays across waves. Overall, much of published cross-sectional results were relevant also in our longitudinal study, indicating that age-associated methylation alterations are persistent over time.

Cellular compositions estimated from methylation data are useful to show how white blood cells change with age. In our longitudinal cohort, T cell compositions were not observed to decrease with age, while the changes of other cell types were in accordance to a previous study (24). The discrepancy in T cells between studies may highlight differences regularly observed between cross-sectional and longitudinal associations where the former sometimes overestimate an association with age. However, it may also be explained by technical artifacts because many samples had cellular estimates very close to zero, making it difficult to perform good regression analyses. In addition, in spite of the observed cellular composition changes with age, they contributed little to the 1316 age-associated CpGs, where estimates of age were similar with and without adjusting for cellular compositions.

Regulatory annotation of age-associated CpGs indicated enrichment in predicted CTCF binding sites, promoter flanking regions and unannotated TF binding sites. DNA methylation has been reported to regulate CTCF binding to DNA, which in turn regulates gene expression through long range interactions with enhancers and promoters (25). Thus, we suggest that this methylation of CTCF is important in the aging process. Hypermethylated CpGs were not enriched in promoter regions, where age-related hypermethylation was believed to occur. Instead, hypermethylated CpGs were commonly associated with TF binding sites than in other regulatory regions, indicating the importance of age-related hypermethylation which occurs in TF binding sites unannotated to known regulatory features. The mechanism of age-related methylation in enhancers and open chromatin regions were not clear, because these regions were poorly covered by the 450k array.

The meQTL analysis identified a large number of CpG-SNP associations and showed that genetic variants have strong effects on DNA methylation. The higher proportion of genetic-associated CpGs found among the age-associated CpGs (9.2%) than in all CpGs (3.7%) indicates enriched genetic impact on age-related methylation. Our results suggested no interaction between genetic and age effects on methylation, which supports the conclusion by Gaunt (12) that genetic effects on methylation are stable over time. Similarly, heritability analyses by Tan (15) showed that intra-individual longitudinal change of age-associated CpGs was mostly (90%) explained by individual specific environmental factors. Thus, genetic variants can affect methylation levels of age-associated CpGs, but not the speed of methylation changes over time. A previous study on methylation heritability (9) suggested gene-age interactions on methylation, but they were not detected in our study.

Furthermore, twin-pair methylation differences using information from all CpGs indicated a global increase in inter-individual methylation variations over time. This increase is probably due to both environmental and stochastic factors. Especially, the increase in methylation differences was stronger (steeper slope) in the age-associated CpGs. Thus, age-associated CpGs not only change with age, but also have higher age-induced variations than average. The result could imply that age-related methylation is more vulnerable to environmental and stochastic effects. Also, comparing MZ and DZ twins showed that genetic effects were stronger in the age-associated CpGs than all CpGs. It corresponded with meQTL results that a higher proportion of SNP-associated CpGs were found in age-associated CpGs.

The strength of this study was the use of longitudinal methylation data with repeated measures sampled up to five times over 20 years in the same individuals. Moreover, we successfully validated our results in two independent longitudinal studies with high correlations of effect sizes. Although blood samples were collected, stored, and DNA extracted at different times, methylation data were measured and processed together with complete randomization. Therefore, we largely reduced the influences of uncorrected batch effects in the analysis. The twin design further enabled us to investigate methylation differences within twin pairs to specifically study environmental influences on methylation.

Among the limitations for this study was that it was performed on a relatively small number of subjects (N=385) and some people only participated in one or two measures. Moreover, the results are only applicable in the old ages and for European ancestry populations.

Further, methylation data were obtained from 450k arrays, which have poor coverage of enhancer regions, an important feature of gene regulation. Also, the quality of data from methylation arrays is not always perfect due to potential unspecific hybridization and noises in signal detection. Stronger evidence could be provided by data from bisulfite pyrosequencing methods as validation.

DNA methylation includes both cell-specific and unspecific patterns. Some evidence show that age-related hypermethylation are more conserved across different tissues than hypomethylation (5, 26). However, we only studied blood samples and adjusted methylation data using estimated cellular compositions (20). Further investigation on cell-type specific and unspecific methylation alterations in longitudinal studies may reveal methylation patterns in relation to cell types, since cellular composition changes across age in blood (24).

## Materials and Methods

### Study aim, design and settings

This study used methylation data of people involved in SATSA (16), which aims to understand individual differences in aging. The SATSA is part of the Swedish Twin Registry (STR) (17), which is a population-based national register including twins born 1886-2000. The SATSA started in 1984, and continuously collected cognitive and health data every third year with up to ten waves in a total of 861 participants.

### Study population

Blood samples were obtained from 402 SATSA participants, including 85 MZ and 116 DZ twin pairs. We collected in total 1122 samples at five time-points starting from 1992 to 2012. After quality control on methylation data, we retained 1011 samples from 385 twins. In the five longitudinal waves, numbers of participants with 1 to 5 measurements were 99, 86, 90, 80, 30 (Table 1). Among them, 200 participants were measured at least three times. Phenotype data were collected through comprehensive questionnaires and physical testing at each sampling wave. Phenotypes used in this study include chronological age, sex, zygosity.

### Methylation data

For each sample, 200 ng of DNA were bisulfite converted using the EZ-96 DNA MagPrep methylation kit (Zymo Research Corp., Orange, CA, USA) according to the manufacturer’s protocol optimized for Illumina’s Infinium 450K assay. The bisulfite converted DNA samples were hybridized to the Infinium HumanMethylation450 BeadChips by the University College London Genomics Core Facility according to Illumina’s Infinium HD protocol (Illumina Inc., San Diego, CA, USA). Samples were randomly distributed into 13 plates. DNA methylation levels of 485,512 CpGs were measured for each sample.

We processed raw methylation data using the R package RnBeads (27). Quality control (QC) was performed in two steps removing; 1) samples having low median signal intensities, wrong predicted sex, or poor correlations (r<0.7) with genetic controls; 2) probes overlapping with a SNP and non-CpG probes. Additionally, we employed a greedy-cut algorithm that iteratively filtered out probes and samples. This was done by maximizing false positive rate minus sensitivity with a detection p-value cutoff of 0.05. In the end, probes on sex chromosomes were removed. After QC, we retained 1011 samples and 390,894 probes.

Subsequently, we used a background correction method “noob” (28) from the methylumi package and a normalization method “dasen” (29) from the watermelon package. Next, we corrected normalized data for cellular compositions, which were estimated by the Houseman method (20) using a blood cell reference panel (30). In order to detect and remove technical variance, we used the Sammon mapping method (31) to achieve a lower-dimension projection preserving the original data structure. The low-dimensional data were then fitted to a linear regression model to test potential batch effects. The strongest batch effects were identified as array slides, and corrected for using the ComBat method from the R package sva (32). Data (beta-values) were then transformed to M-values for statistical analysis through a logit transformation.

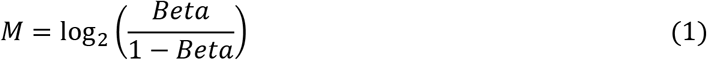

### Genotype data and imputation

We generated genotype data using the Illumina PsychChip (Illumina Inc., San Diego, CA, USA), which detected 588,454 SNPs for each individual. We applied QC criteria by removing; 1) samples having >1% missing genotype, estimated inbreeding coefficient >3 standard deviations from the sample mean, wrong relatedness between individuals and wrong predicted sex; 2) SNPs not mapped to a chromosome, with over 2% missing calls, deviating from the Hardy-Weinberg equilibrium (p<10^−6^) and with no observed minor alleles.

After QC, we performed pre-phasing on genotype data using SHAPEIT v2.r837 with default parameters. Imputation was then performed in chunks of around 5 Mb using IMPUTE2 version 2.3.2 with default parameters (33). The imputation reference was based on the 1000 Genomes Project phase 1 (34). Next, a QC step was performed on imputed genotype data to filter out low imputation quality variants (Info<0.6) and low minor allele frequency variants (MAF < 0.05). After imputation and QC, in total 363 individuals from SATSA had genetic data including 6,528,198 imputed SNPs.

### PIVUS and LBC cohorts

The PIVUS (18) study included 390 samples from 196 individuals collected at two specific ages, 70 and 80 years. Half of the participants from PIVUS were women. Methylation data were obtained from blood samples using the Illumina 450k array which has been described previously (35).

The LBC study was composed of two different birth cohorts, LBC1936 and LBC1921 (19), with 3018 samples collected in max of three time points. In the three waves, LBC1936 included 906, 801 and 619 individuals with mean ages of 69.6, 72.5 and 76.3 years; LBC1921 included 436, 174, 82 individuals with mean ages of 79.1, 86.7 and 90.2 years. Proportions of women were 49.4% and 53.7% in LBC1936 and LBC1921 at baseline. Methylation data were obtained from blood samples using the Illumina 450k array as presented elsewhere (13).

### Statistical analyses

We fitted a linear mixed model to describe longitudinal changes of methylation with age. The model included fixed effects of age and sex, and random intercepts and slopes between twins nested in twin pairs. In the model formula below, *i, j* and *k* denote twins, twin pairs and time points; *γ, β_1_, β_2_, u, ω* and *ε* denote fixed intercept, fixed coefficient of age, fixed coefficient of sex, random intercept, random coefficient of age and random error.

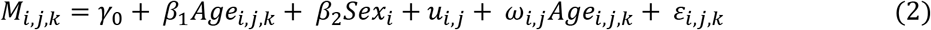

The sensitivity analysis on twin zygosity adjusted zygosity as a random effect in the mixed model.

We used two longitudinal cohorts, PIVUS and LBC, to validate our results. In PIVUS validation, a linear regression model using general least squares was performed to estimate methylation changes over 10 years for each CpG. Confounders included sex, smoking status, cell counts and batch effects were adjusted for in the model. In LBC validation, a mixed effect model allowing for random intercepts and slopes of age were performed to measure age-associated methylation longitudinally. Cell counts was adjusted as a covariate, and sex, sub-cohort, plates, array, position and hybridization date were adjusted as random effects.

The *cis*-meQTL analysis was performed on 363 participants in SATSA with both genotype and methylation data available. Methylation data from the first observation were used to identify meQTLs. To reduce the computational complexity, we employed the R package matrixEQTL (36) to perform a fast rough screening for *cis*-meQTLs. Genotypes were treated to have additive effects in the model. The screening method used a linear regression model, including age, sex and the first four genetic principle components as covariates, to calculate all *cis*-methylation-genotype associations. We selected associations with p<1×10^−8^ from the screening results and further fit to a linear regression model including the same covariates. After that, sandwich estimators were used to correct standard errors for the effect of having correlated observations from twins in the sample.

The intra-twin-pair differences of methylation patterns were measured by Euclidean distances. Methylation distances were calculated within twin pairs at the same time point across the CpGs used. In total 154 complete twin pairs (69 MZ, 85 DZ pairs, 660 samples) were available at the same time point. Methylation distances were standardized before regression.

Three sets of methylation distances were calculated using all CpGs, age-associated CpGs and SNP-associated CpGs respectively. Then, a linear mixed effect model was fitted to measure how intra-twin-pair methylation differences changed over time. The mixed effect model included age, sex and twin zygosity as fixed effects and twin pair as the random effect shown below, where *i* and *j* denote twin pairs and time points, *y, β_i_, β_2_, β_3_, u*, and *ε* denote fixed intercept, fixed coefficient of age, fixed coefficient of sex, fixed coefficient of zygosity, random intercept and random error.

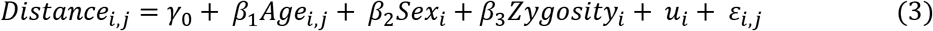

The probe-wise methylation distances were calculated from the absolute difference between M-values. The same regression model (Equation 3) was employed to estimate the change of intra-twin-pair distances overtime.

### Regulatory annotation and functional analysis

We annotated genome locations and related regulatory features of identified age-associated CpGs using Ensembl Funcgen database (21). Regulatory features were classified according to the Ensembl Regulatory Build (37). In addition, we performed functional annotation on enriched genes using DAVID (22, 23) online tools.

### Ethics approval and consent to participate

All participants in SATSA, PIVUS and LBC have provided written informed consents. This study was approved by the ethics committee at Karolinska Institutet with Dnr 2015/1729-31/5.

### Availability of data and material

The datasets generated and/or analysed during the current study are available in the Gene Expression Omnibus repository, [PERSISTENT WEB LINK TO DATASETS]

### Competing interests

EI is a scientific advisor for Precision Wellness, Cellink and Olink Proteomics for work unrelated to the present project.

## Funding and Acknowledgements

The SATSA study was supported by NIH grants R01 AG04563, AG10175, AG028555, the MacArthur Foundation Research Network on Successful Aging, the Swedish Council for Working Life and Social Research (FAS/FORTE) (97:0147:1B, 2009-0795, 2013-2292), the Swedish Research Council (825-2007-7460, 825-2009-6141, 521-2013-8689, 2015-03255), Karolinska Institutet delfinansiering (KID) grant for doctoral students (YW), the KI Foundation, and by Erik Rönnbergs donation for scientific studies in aging and age-related diseases.

